# Extra-islet expression of islet antigen boosts T-cell exhaustion to prevent autoimmune diabetes

**DOI:** 10.1101/2023.02.12.528226

**Authors:** Claudia Selck, Gaurang Jhala, David De George, Chun-Ting J. Kwong, Marie K. Christensen, Evan Pappas, Xin Liu, Tingting Ge, Prerak Trivedi, Axel Kallies, Helen E. Thomas, Thomas W.H. Kay, Balasubramanian Krishnamurthy

## Abstract

Persistent antigen exposure results in the differentiation of functionally impaired, also termed exhausted, T cells which are maintained by a distinct population of precursors of exhausted T (T_PEX_) cells. T cell exhaustion is well studied in the context of chronic viral infections and cancer, but it is unclear if and how antigen-driven T cell exhaustion controls progression of autoimmune diabetes and whether this process can be harnessed to prevent diabetes. Using non-obese diabetic (NOD) mice, we show that some CD8+ T cells specific for the islet antigen, islet-specific glucose-6-phosphatase catalytic subunit-related protein (IGRP) displayed terminal exhaustion characteristics within pancreatic islets but were maintained in the T_PEX_ cell state in peripheral lymphoid organs. To examine the impact of antigen on T cell exhaustion in diabetes, we generated transgenic NOD mice with inducible IGRP expression in peripheral antigen presenting cells. Antigen exposure in the extra-islet environment induced severely exhausted IGRP-specific T cells with reduced ability to produce IFNγ, which protected these mice from diabetes. Our data demonstrate that T cell exhaustion induced by delivery of antigen can be harnessed to prevent autoimmune diabetes.

## Introduction

Chronic antigen exposure in cancer and chronic viral infection causes CD8^+^ T cells to undergo progressive cellular differentiation with loss of cytokine production and cytotoxicity, a state termed T-cell exhaustion (1–3). T cell exhaustion is a physiological adaptation to continuous antigen stimulation that protects against excessive immune-mediated tissue damage. However, it also results in failure to clear the antigen leading to viral or tumor persistence. T cell exhaustion has also been described in autoimmunity driven by immune mediated tissue damage (4). In type 1 diabetes (T1D), CD8^+^ T cell mediated beta-cell destruction takes years in humans and months in the non-obese diabetic (NOD) mouse model (5–7). This chronicity is likely to be due to a balance between the autoimmune attack and processes such as T cell exhaustion, that reduce its effectiveness. This balance is complex and at present insufficiently understood.

Exhausted T cells are heterogenous, consisting of subsets with distinct functional properties. Exhausted T cells are maintained by precursors of exhausted T (T_PEX_) cells (8–10). These cells, which retain high proliferative potential and undergo long-term self-renewal, express the transcription factor TCF1 (encoded by *TCF7*). They give rise to short-lived terminally exhausted effector T (T_EX_) cells with restrained functionality that express TIM-3 but lack TCF1 (11). T_PEX_ cells mediate the response to therapies that block the immune checkpoint programmed cell death 1 (PD-1) and thereby reinvigorate exhausted T cells in cancer (8–10). TCF1+ exhausted T cells with precursor properties have also been described from a range of autoimmune conditions (12–14). Transcriptomic profiling of T cells from patients with other conditions, such as vasculitis, Crohn’s disease and systemic lupus erythematosus, showed that a T cell exhaustion signature correlated with a more benign form of autoimmune disease, indicating that mechanisms associated with T-cell exhaustion may be important in controlling autoimmunity (15). Consequently, PD-1 blocking therapies can precipitate or exacerbate autoimmunity (16–18).

The T cell response to islet-specific glucose-6-phosphatase catalytic subunit-related protein (IGRP), a major autoantigen recognised by CD8^+^ T cell in NOD mice(19, 20) allows the study of CD8^+^ T cell exhaustion in spontaneously developing polyclonal antigen-specific T cells in a clinically relevant model of autoimmunity. Recent studies have identified exhausted CD8^+^ T cells in islets of NOD mice (12, 21, 22), although one study (23) suggested that CD8^+^ T cells do not display transcriptional or phenotypic exhaustion in islets of NOD mice. Transcriptional profiling has identified T_PEX_ and T_EX_ subsets of CD8^+^ T cells that infiltrate the pancreatic islets of NOD mice; these exhausted CD8^+^ T cells expand as mice age (22). Progression of type 1 diabetes is slower in individuals with islet-specific CD8^+^ T cells displaying an exhausted phenotype with expression of multiple inhibitory receptors (24). While reinvigoration of exhausted CD8^+^ T cells with checkpoint inhibitors is the aim of cancer treatment, we reasoned that driving terminal differentiation of exhausted CD8^+^ T cells may be a strategy to prevent diabetes in NOD mice. T_PEX_ reside in lymphoid tissues. When these cells encounter antigen they proliferate, leave the lymph nodes and enter the peripheral tissues via the blood and during this process they lose TCF1 expression and differentiate to T_EX_ CD8^+^ T cells. We hypothesised that CD8^+^ T cell exhaustion could be boosted by continuously exposing T cells to antigen while resident in lymph nodes. To test this idea, we generated tetracycline-inducible IGRP transgenic non-obese diabetic (NOD) mice to enable temporal expression of IGRP in antigen presenting cells (APCs) in a doxycycline dependent manner (25). Antigen expressed in the peripheral lymphoid organs induced terminal exhaustion in CD8^+^ T cells that subsequently migrated to the islets. Surprisingly, while deletion of CD8^+^ T cells specific for IGRP_206–214_ (IGRP-specific T cells) did not prevent diabetes, boosting exhaustion in IGRP-specific T cells prevented diabetes in NOD mice. Overall, we show that boosting T cell exhaustion in autoimmune diabetes is a promising strategy to limit the progression of disease.

## Results

### Islet antigen specific T cells undergo an exhaustion program in NOD mice

Several studies described features of exhaustion for islet-infiltrating CD8^+^ T cells in NOD mice (12, 21, 22). Furthermore, a recent study showed that IGRP-specific T cells in the draining pancreatic lymph node expressed high levels of TCF-1 and gave rise to functional effector T cells, which infiltrated the pancreas and destroyed insulin producing beta cells (23). This suggested that IGRP-specific T cells undergo only partial exhaustion and retain effector function. To better understand T-cell exhaustion and heterogeneity, we performed single cell RNA-seq analysis of immune cells in the islets (Supplementary Fig S1). Consistent with previous analyses, the infiltrate was dominated by T cells, including CD4 and CD8 T cells, and B cells with smaller populations of NK cells and myeloid cells (Supplementary Fig S1A-D). Subsequent analyses, restricted to antigen-responsive (CD44high) T cells (Fig 1A), identified exhausted CD8^+^ T cells expressing *Tox* and *Pdcd1* (encoding PD-1) (Fig 1B). These cells could be further segregated into T_PEX_ cells expressing *Tcf7, Slamf6* and *Eomes*, associated with self-renewal and memory function, and T_EX_ cells that were negative for *Tcf7* but expressed a range of transcripts encoding co-inhibitory receptors and transcription factors associated with terminal exhaustion, including TIM-3 (*Havcr2*), Lag3, Id2 and Blimp1 (*Prdm1*) (Fig 1B, C). Notably, among the exhausted T cells, T_PEX_ cells were the predominant population (Fig 1A, B), which was surprising as in chronic viral infection and in tumors T_PEX_ cells are predominantly found in lymphoid tissues, while Tex cells preferentially reside in non-lymphoid tissues (8).

**Figure 1:**
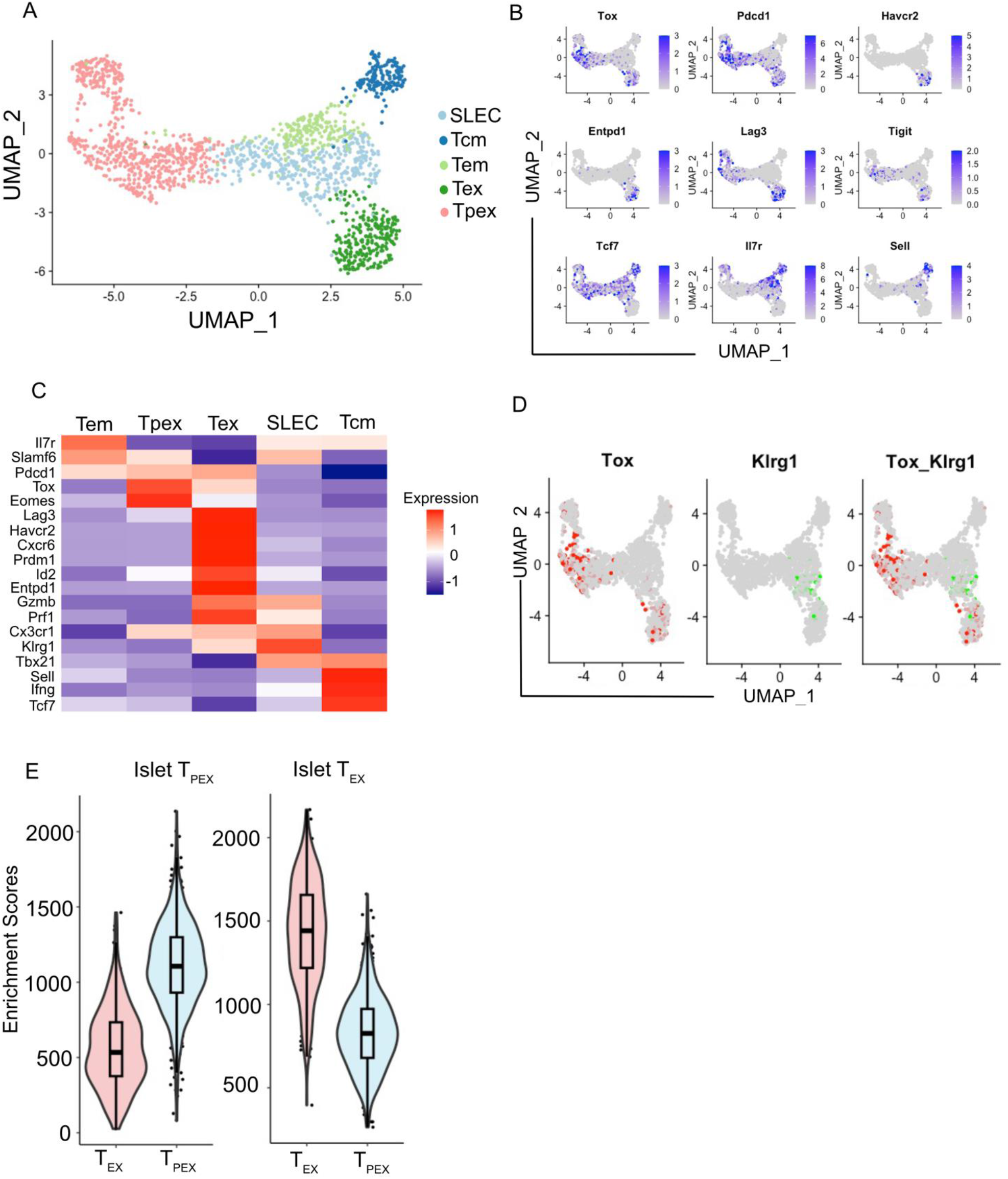
Islet antigen specific T cells undergo an exhaustion program in NOD mice. (A) Uniform Manifold Approximation and Projection (UMAP) plot of islet infiltrating CD44+ CD8+ T cells in 16-week-old NOD mice showing short lived effector (SLEC), central memory (Tcm), effector memory (Tem), precursor of exhausted (T_PEX_) and terminal exhausted (T_EX_) subsets of CD8+ T cells. (B) Expression of indicated genes in individual cells from A. (C) Heatmap of scRNA-seq showing row-normalized log2 (counts per million) for differentially expressed genes in T_EX_, T_PEX_, short lived effector cells (SLEC) and memory (Mem) CD8+ T cells in islets of NOD mice. (D) Expression of klrg-1 and Tox genes in individual cells from A. (E) Gene set enrichment analysis of a signature of islet infiltrating T_PEX_ and T_EX_ CD8+ T cells from NOD mice using a core set of genes differentially expressed between T_PEX_ and T_EX_ in both chronic LCMV infection and the B16-OVA model of tumour immunity.

In addition to exhausted T cells, we also found T cells with memory characteristics that lacked expression of *Tox* and transcripts of co-inhibitory receptors. This included central memory cells expressing *Tcf7, Il7r* and *Sell* (encoding CD62L), and an effector memory subset lacking expression of *Sell* (Fig 1A-C). Furthermore, we identified a cluster that was enriched for *Klrg1* expressing cells (short-lived effector cells, SLEC), which did not overlap with Tox expressing exhausted cells (Fig 1A-D). Genes associated with cytotoxicity, including Gzmb and prf1 (encoding perforin), were expressed in both the T_EX_ cell and SLEC clusters. Gene set enrichment analysis using a core set of genes differentially expressed between T_PEX_ and T_EX_ in both chronic LCMV infection and the B16-OVA model of tumour immunity (11) showed that both our T_PEX_ and T_EX_ cell clusters were enriched for expression of genes upregulated in classical T_PEX_ and T_EX_ populations in chronic infection and cancer (Fig 1E). Thus, T cells that have entered an exhaustion program comprise a large fraction of islet infiltrating CD8 T cells.

### IGRP-specific T cells undergo exhaustion but are maintained as T_PEX_ cells in NOD mice

We next analyzed IGRP-specific T cells from islets, pancreatic lymph nodes (PLN) and peripheral lymphoid organs (PLO, inguinal and mesenteric lymph nodes and spleen) of NOD mice by magnetic bead enrichment and tetramer staining (Supplementary Fig S2). We found that all the IGRP-specific T cells, whether isolated from islets, pancreatic lymph nodes or PLO, expressed TOX and could thus be classified as exhausted T cells. The expression of TOX was highest in IGRP-specific T cells isolated from the islets followed by PLN and PLO (Fig 2A). We detected high levels of the inhibitory receptors PD-1, TIM-3 and TIGIT on islet-infiltrating IGRP-specific T cells (Fig 2B, C). About 60-80% of the IGRP-specific T cells in the islets displayed a Slamf6^+^TIM-3^-^ T_PEX_ cell phenotype (Fig 2D), while the remaining 20-40% of the cells were TIM-3^hi^ T_EX_ cells. As expected, Slamf6 and TCF-1 were co-expressed (Supplementary Fig S3A).

**Figure 2:**
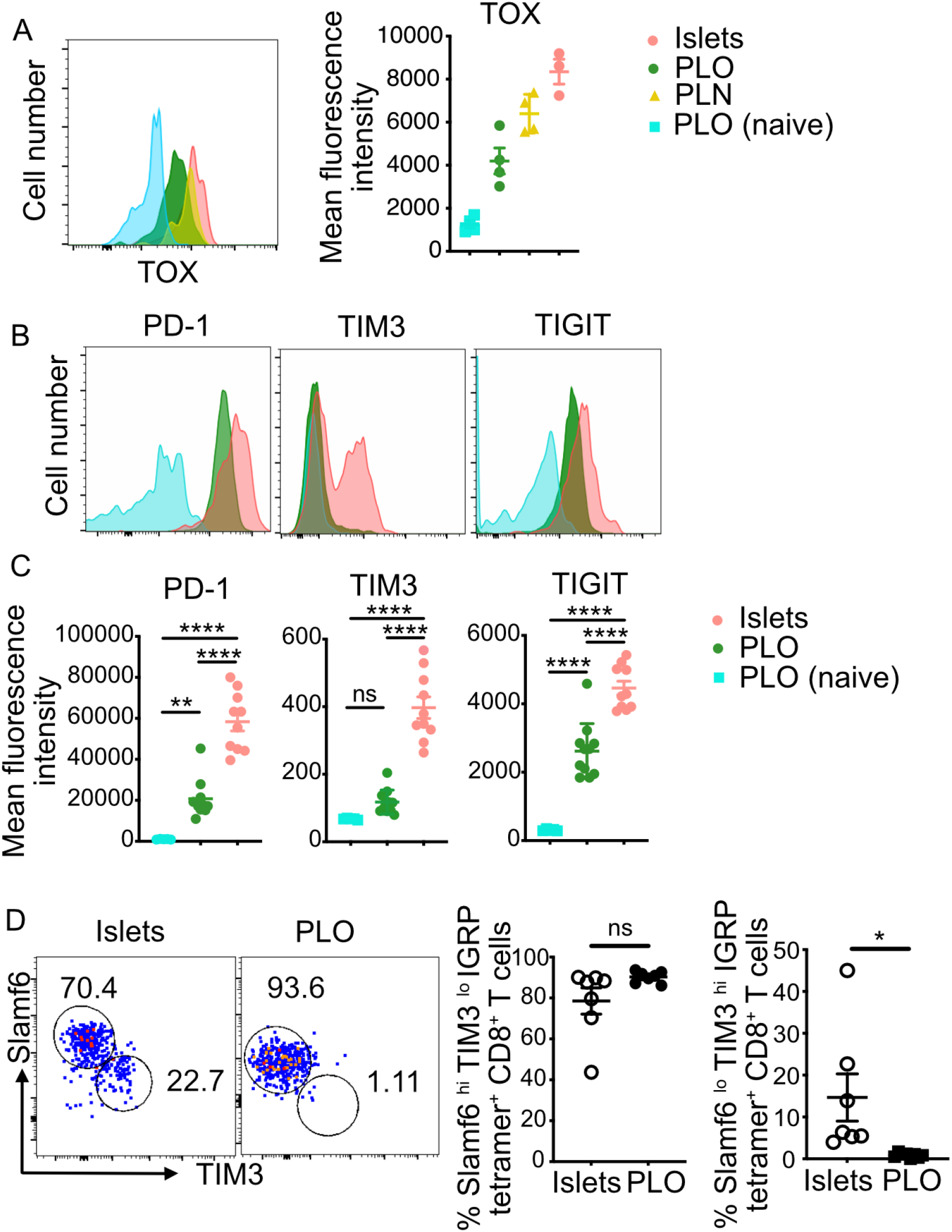
Precursor of exhausted and terminal exhausted antigen-specific T cells are found in islets of NOD mice. IGRP-specific T cells and naïve CD8+ T cells from islets and peripheral lymphoid organs (PLO, non-draining lymph nodes and spleen) of 16-18-week-old NOD mice were analysed by flow cytometry after tetramer staining and magnetic bead enrichment. (A) Representative histogram and mean fluorescence intensity of TOX expression in IGRP-specific T cells from islets, PLN and PLO as compared to naïve CD8+ T cells in the PLO. (B) Representative histograms showing the expression of indicated co-inhibitory receptors and (C) quantification of mean fluorescence intensity (MFI) in the islets and PLO of NOD mice. (D) Proportion of Slamf6^hi^TIM3^lo^ precursor of exhausted and Slamf6^lo^TIM3^hi^ terminal exhausted IGRP-specific CD8+ T cells from PLO and islets of 16-18-week-old NOD mice. Pooled data (mean ± SEM) in (B-D) from 3 independent experiments, each symbol in the scatter plots represents data from an individual mouse. Values in the FACS plots (D) show percentages. ***P < 0.001, **** P < 0.0001, using 2-tailed unpaired t test.

In the draining pancreatic lymph nodes, IGRP-specific T cells were TOx^hi^PD-1^hi^, indicating that they had undergone an exhaustion program. While the majority expressed Slamf-6 and TCF-1, only about 5-10% expressed TIM-3 (Supplementary Fig S3B), indicating that IGRP-specific T cells in the PLN were mainly T_PEX_ cells. Consistent with our previous observation which showed that antigen-experienced IGRP-specific T cells in the PLO of NOD mice are derived from islet-infiltrating cells (26), IGRP-specific T cells in other lymphoid organs resembled the phenotype of T cells in the PLN. While the expression of PD-1 was lower on IGRP-specific T cells in the PLO compared to the islets (Fig 2B, C) it was greater than on naïve or CD44^hi^, KLRG-1^hi^ effector CD8^+^ T cells not specific for IGRP (Supplementary Fig S3C), indicating that exhaustion features were imprinted even when the cells leave the site of antigen exposure in the islets. Thus islet-specific T cells in NOD mice do undergo an exhaustion program within the islets but are predominantly maintained as self-renewing T_PEX_ cells in the islets and recirculate in the PLO.

### Temporal expression of IGRP in the antigen-presenting cells of NOD mice

To directly study the impact of antigen exposure on IGRP-specific T cells we made use of transgenic NOD mice in which IGRP is expressed under the control of the MHCII (I-Eακ) promoter and the expression of IGRP in antigen presenting cells (APCs) is prevented by exposure to doxycycline (25) (Fig 3A), termed <u>T</u>etracycline <u>I</u>nhibited <u>I</u>GRP (<u>NOD-TII) mice</u>. To examine the fidelity and robustness of IGRP transgene expression, NOD-TII mice were induced to express IGRP at 10 weeks of age. At 12, 14 and 16 weeks, spleens were harvested for longitudinal analysis of transgenic IGRP expression via qPCR (Fig 3B, C). These were compared to NOD-TII mice that expressed IGRP from birth (never given doxycycline, these mice were called NOD IGRP mice) and those that never expressed IGRP (doxycycline from gestation). There was no difference in IGRP expression in the APCs from spleens of 10-week-old transgenic NOD mice that were never induced to express IGRP and wildtype NOD mice, indicating that the IGRP expression was not ‘leaky’ in the transgenic NOD-TII mice (Supplementary Fig S3D). From here on this cohort is called ‘NOD control’. After cessation of doxycycline at 10 weeks of age, IGRP expression was significantly higher in 12-week-old NOD-TII mice compared with NOD control mice, and this transcriptional level was maintained (Fig 3C). However, this expression was lower than when IGRP was induced from birth (NOD-IGRP mice, Fig 3C). To confirm IGRP antigen expression, we CFSE-labelled T cells from NOD8.3 TCR transgenic mice that have ~90% of CD8^+^ T cells specific for IGRP_206–214_ and transferred them into NOD-TII and NOD control mice. As expected, in NOD control mice, transferred NOD8.3 T cells proliferated only in the islets where native IGRP is found and in the draining PLN (Fig 3D). In contrast, extensive proliferation of the transferred NOD8.3 T cells in NOD-TII mice was detected in both the PLN and non-draining lymph nodes confirming that IGRP is expressed by the APCs of NOD-TII mice.

**Figure 3:**
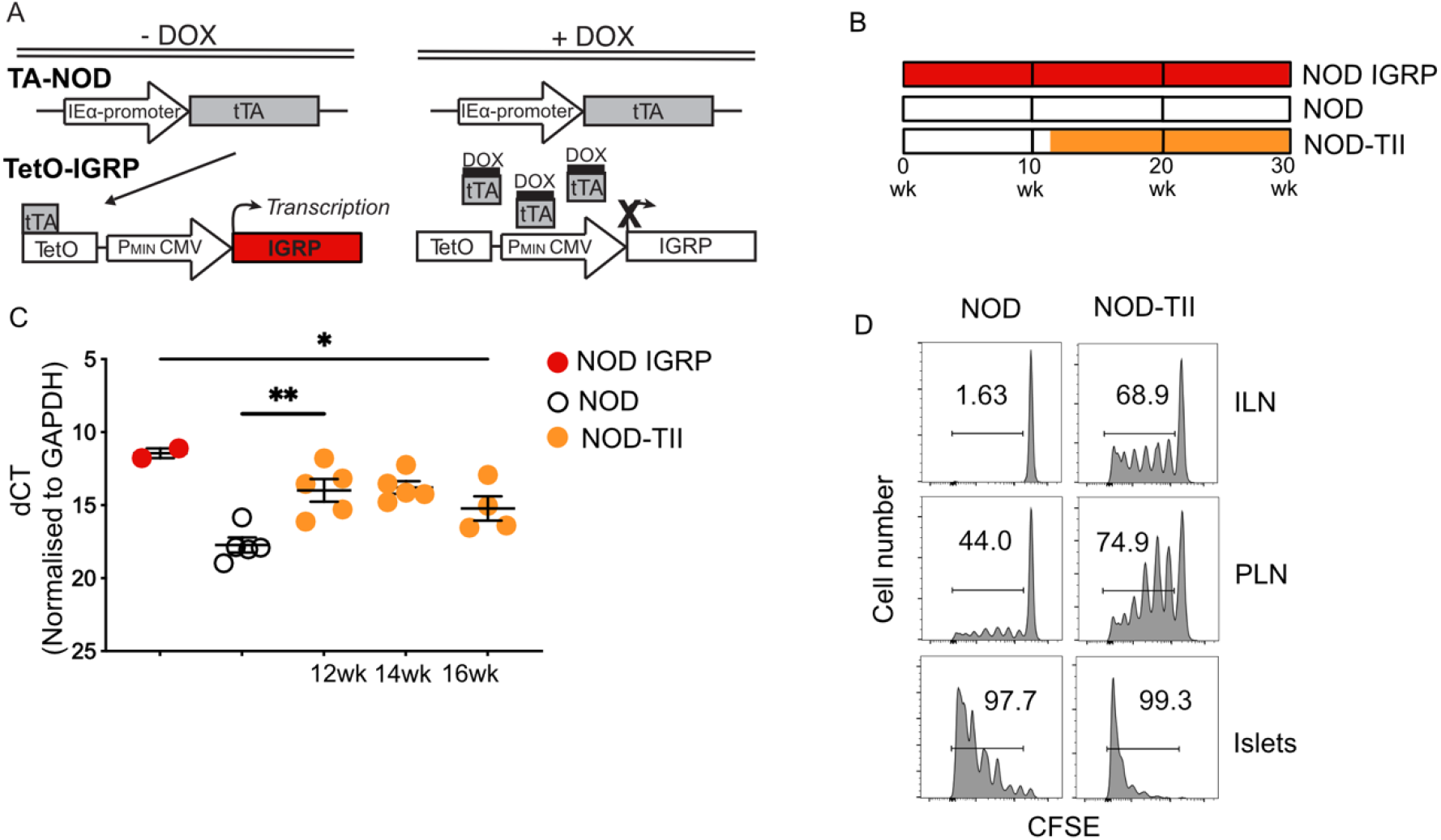
Doxycycline regulated IGRP expression in NOD-TII mice. (A) Scheme of generation of tetracycline regulated NOD-IEα-tTA (TA-NOD) and TetO-IGRP dual transgenic mice (NOD-TII mice). (B) Study design depicting treatment groups. Continuous IGRP expression in the absence of doxycycline (Dox) shown in red (NOD.IGRP mice), no IGRP expression while always on Dox in white (control NOD mice) and IGRP expression induced after removal of Dox at 10 weeks of age shown in orange (NOD-TII mice). (C) Quantitative RT-PCR for *G6pc2* in splenic lysates of NOD.IGRP, NOD and NOD-TII mice at indicated ages. Data show dCT values (mean±SEM) of individual mice from 2 independent experiments. * p<0.05, **p<0.01, one-way ANOVA with Tukey’s multiple comparisons test. (D) CFSE labelled CD8^+^ T cells from NOD8.3 mice were transferred into NOD recipients at 10-12 weeks of age and into NOD-TII recipients 6 weeks after stopping Dox treatment to induce IGRP expression. CFSE labelled cells were analyzed in the islets, pancreatic lymph nodes (PLN) and inguinal lymph nodes (ILN) 5 days post-transfer. Data are representative of 2 independent experiments. Numbers in each histogram indicate percentage of CFSE low cells.

There was no difference in proliferation of transferred cells in the islets of NOD control or NOD-TII mice indicating that there is already abundant endogenous IGRP in the islets. The extent of proliferation of transferred NOD8.3 T cells in inguinal lymph nodes of NOD-TII mice was less than that seen in islets of either NOD control or NOD-TII mice, indicating that IGRP specific T cells were stimulated less in the lymph nodes than in islets of NOD-TII mice (Fig 3D). This was likely to be due to the lower expression of costimulatory markers CD86, CD40 and MHC class II on APC from PLO compared to APCs from islets (Supplementary Fig S3E). Together, these observations confirm the fidelity and robustness of NOD-TII transgenic mice.

### Autoantigen expression in the periphery induces functional T cell exhaustion

As we have previously described (20), IGRP-specific T cells in transgenic NOD-TII mice expressing IGRP from birth were almost completely deleted (Fig 4A, NOD-IGRP). We next induced IGRP expression from 10 weeks of age. Notably, antigen-experienced CD44^hi^ IGRP-specific T cells in the PLO were detectable at similar frequencies in 20-week-old NOD-TII mice (induced to express IGRP from 10 weeks of age) and NOD control mice (never induced to express IGRP) (Fig 4A, B), indicating that, unlike naïve T cells, antigen-experienced IGRP-specific T cells are refractory to deletional tolerance upon antigen exposure.

**Figure 4:**
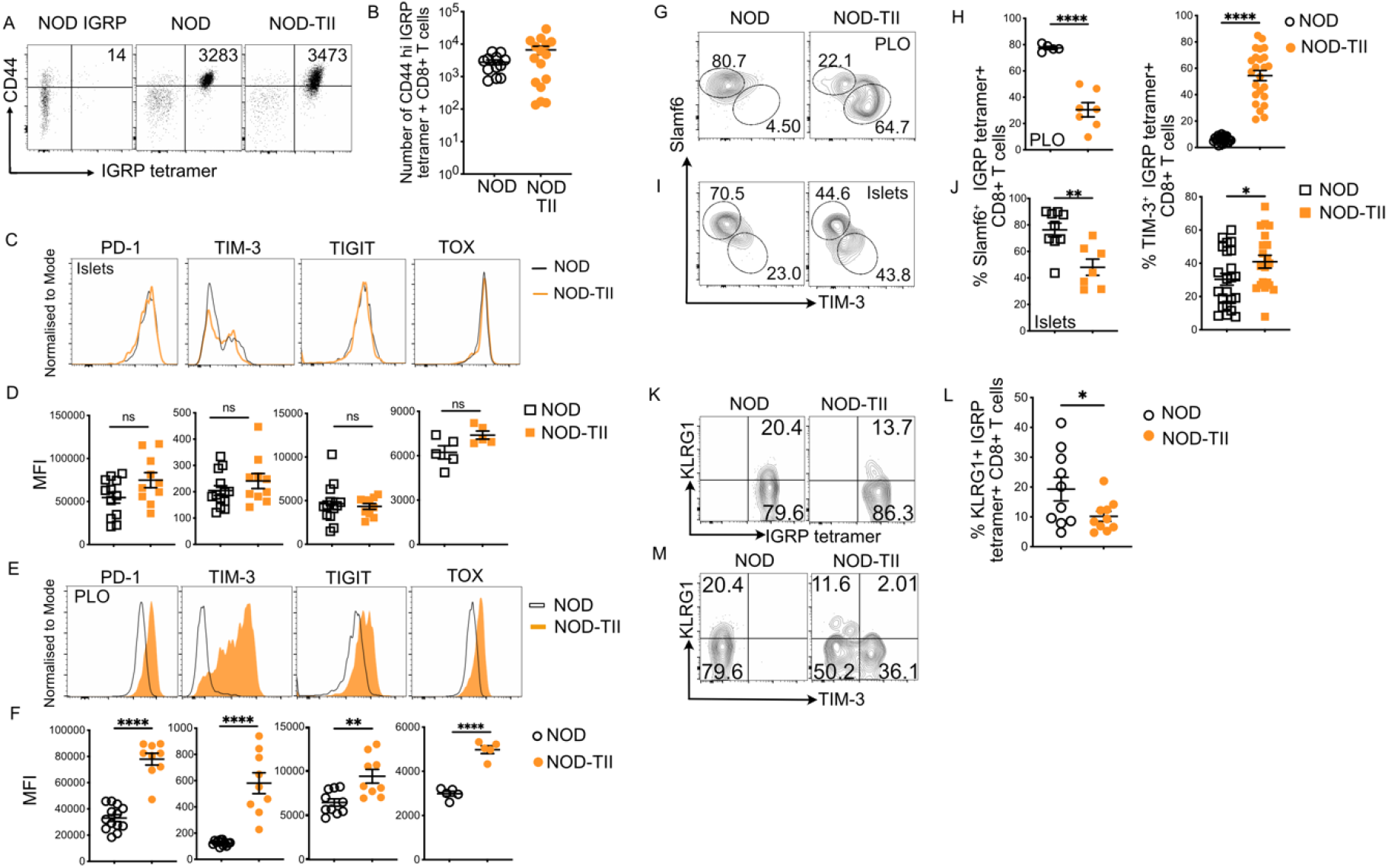
Ongoing antigen exposure induces markers of T-cell exhaustion on IGRP-specific T cells. IGRP-specific T cells were stained with K^d^-IGRP_206–214_ tetramer and enriched from pooled peripheral lymphoid organs (PLO) of 16–18-week-old NOD and NOD-TII mice using magnetic beads and enumerated by flow-cytometry. IGRP-specific T cells were analysed in dispersed islets by tetramer staining without enrichment. (A) Representative FACS plots and (B) quantification of absolute number of IGRP-specific CD8^+^ CD44^hi^ T cells after enrichment from PLO. Values in the FACS plots indicate absolute number of tetramer binding cells. (C) Representative histograms and (D) mean fluorescence intensity (MFI) quantification of PD-1, TIM-3, TIGIT and TOX on IGRP-specific T cells in islets of indicated mice. (E) Representative histograms and (F) mean fluorescence intensity (MFI) quantification of PD-1, TIM-3, TIGIT and TOX on IGRP-specific T cells in PLO of indicated mice. (G) Representative FACS plots and (H) frequency of Slamf6 and TIM-3 expressing IGRP-specific T cells in PLO of NOD and NOD-TII mice. Numbers in (G) are the frequency of cells in each circled population. (I) Representative FACS plots and (J) frequency of Slamf6 and TIM-3 expressing IGRP-specific T cells in islets of NOD and NOD-TII mice. Numbers in (I) are the frequency of cells in each circled population. (K) Representative FACS plots and (L) frequency of KLRG-1 expressing IGRP-specific T cells in PLO of NOD and NOD-TII mice. (M) Representative FACS plots frequency of KLRG-1 expressing cells in TIM-3^+^ and TIM-3^-^ IGRP-specific T cells in PLO of NOD and NOD-TII mice.

Islet infiltrating IGRP-specific T cells in both NOD-TII and NOD control mice uniformly expressed high levels of TOX, PD-1, TIM-3 and TIGIT (Fig 4C, D). IGRP-specific T cells in the PLO of NOD-TII mice expressed higher levels of TOX, PD-1, TIM-3 and TIGIT compared to NOD control mice (Fig 4E, F). We next compared the subsets of exhausted IGRP-specific T cells from islets of NOD control and NOD-TII mice (Fig 4I, J). Substantially larger proportions of IGRP-specific T cells were T_EX_ cells in the PLO (Fig 4G, H) and islets (Fig 4I, J) of NOD-TII mice compared to NOD control mice (Fig 4G, H, I, J). Indeed in PLO, while almost all IGRP-specific T cells in NOD control mice were Slamf6^+^TIM-3^-^ T_PEX_ cells, only about 30% were T_PEX_ cells in NOD-TII mice (Fig 4G, H). Also, while a subset of IGRP specific T cells from PLO of NOD mice expressed KLRG-1, this was significantly decreased in NOD TII mice (Fig 4K, L) and KLRG-1 was predominantly expressed by the T_PEX_ subset of IGRP specific T cells in both NOD and NOD TII mice (Fig 4M). These results indicate that continuous exposure to autoantigen in the periphery drives T_EX_ cell differentiation.

### IGRP-specific T cells are functionally disabled when exposed to antigen and recover after termination of antigen exposure

We next assessed the function of IGRP specific T cells in NOD TII mice. The pool of TIM-3^+^ T_EX_ cells can be further dissected based on the expression of the chemokine receptor CX3CR1. While CX3CR1+ cells maintain the highest effector function, loss of CX3CR1 demarcates acquisition of a terminally exhausted state with low effector function and survival (27–29). About 40 % of TIM-3^+^ IGRP-specific T cells in the islets expressed CX3CR1 and there was no difference between NOD control and NOD-TII mice (Fig 5A). However, in the PLO of NOD control mice the majority of TIM3^+^ cells expressed CX3CR1 while only about 20% of the cells expressed CX3CR1 in NOD-TII mice (Fig 5A, B). Consistent with this observation, IGRP specific cells from the PLO of NOD-TII mice expressed less IFNγ following stimulation with PMA and Ionomycin *in vitro* compared with NOD mice (Fig 5C, D). Although the majority of IGRP specific cells in NOD-TII mice displayed phenotypic markers of terminal exhaustion and decreased IFN-γ secretion, the number of IGRP specific cells was not decreased suggesting that the self-renewal and precursor function of T_PEX_ cell remained intact despite high antigen load.

**Figure 5:**
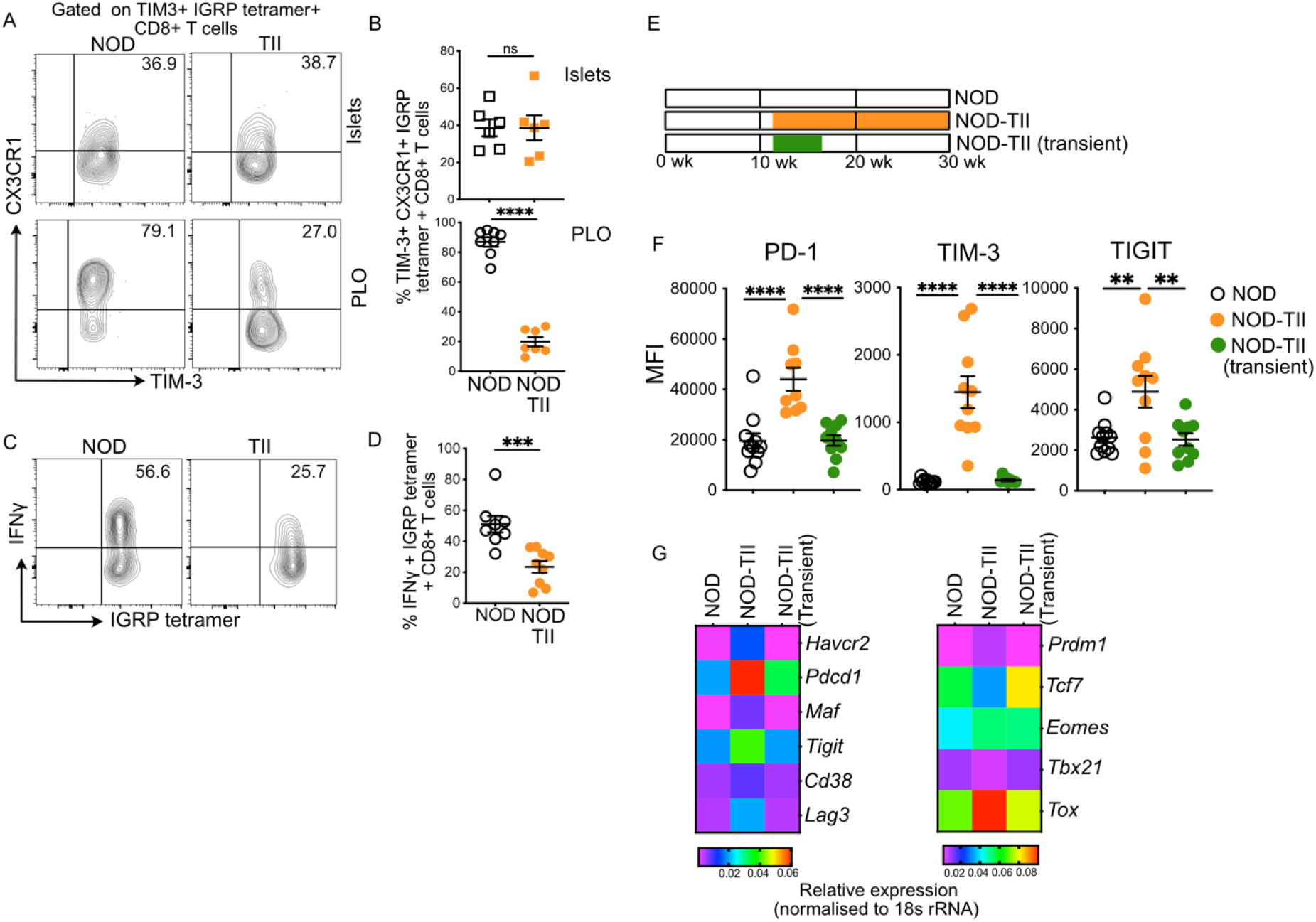
IGRP-specific T cells are functionally disabled when exposed to antigen and recover after termination of antigen exposure. (A) Representative FACS plots and (B) frequency of CX3CR1 and TIM-3 expressing IGRP-specific T cells in islets (top panel) and enriched from PLO (bottom panel) of NOD and NOD-TII mice. The panels A and B show cells gated on Tim-3+, IGRP-specific T cells. Numbers in (A) are the frequency of CX3CR1 expressing cells. (C) Representative FACS plots and (D) frequency of IFNγ expressing IGRP-specific T cells analysed after re-stimulation with PMA and ionomycin for 4 hours. Numbers in (C) are the frequency of IFNγ expressing cells. Data in the scatter plots show mean±SEM from individual mice, pooled from 3-4 independent experiments. Statistical analysis performed using unpaired t-test. ns= not significant, * p<0.05, **p<0.01, ***p<0.001, ****p<0.0001. (E) Study design depicting treatment groups. NOD never expressed IGRP (white), NOD-TII expressed IGRP from 12 weeks of age (orange) and NOD-TII transient expressed IGRP from 16-20 weeks of age (green). (F) MFI quantification of PD-1, TIM-3 and TIGIT on IGRP-specific T cells enriched from PLO of NOD, NOD-TII and NOD-TII mice with transient IGRP expression at 20 weeks of age, analysed by flow-cytometry. (G) Expression of exhaustion-associated genes on IGRP-specific T cells sorted from PLO of NOD, NOD-TII and NOD-TII mice with transient IGRP expression, assessed by quantitative RT-PCR. Heatmaps showing relative expression normalised to 18srRNA. Data pooled from 2-3 independent experiments. Statistical analysis performed using one-way ANOVA with Tukey’s post-test (B-D) or unpaired t-test (F). ns= not significant, * p<0.05, **p<0.01, ***p<0.001, ****p<0.0001.

To examine the stability of phenotypic exhaustion in polyclonal IGRP-specific cells, we induced IGRP expression in NOD-TII mice only for a defined period, between 10 and 15 weeks of age (Fig 5E). As above, T-cell exhaustion markers were upregulated at 15 weeks of age. Following 5 weeks without antigen expression (at 20 weeks of age), IGRP specific cells in the PLO had downregulated expression of inhibitory receptors PD-1, TIGIT and TIM-3 (Fig 5F) to a level that was not different to IGRP specific cells from NOD control mice never exposed to transgenic expression of IGRP. There was also no difference in the number of IGRP specific cells in the PLO (Supplementary Fig S3F) or their ability to secrete IFNγ following stimulation with PMA and ionomycin in vitro (Supplementary Fig S3G) at 20 weeks. We next sorted IGRP-specific cells from the PLO of NOD-TII mice and performed RT-PCR for the expression of exhaustion marker genes. The expression of exhaustion related genes (*Pdcd1*, *Tox, Havcr, Prdm1, Maf, Cd38(30), Lag3* and *Tigit*) was similar in IGRP-specific cells from NOD-TII mice after transient transgene expression and NOD control mice, while it was increased in NOD-TII mice with continuing transgene expression (Fig 5G). *Tcf7, Eomes*, and *Tbx21* were decreased in NOD-TII mice with current antigen expression and recovered after removal of antigen to the level seen in NOD-TII mice that never expressed antigen (Fig 5G). Overall, these results demonstrate that T_PEX_ cells remain functional during periods of high antigen expression and mediate recovery of auto-aggressive IGRP-specific cells following antigen withdrawal.

### Exhausted IGRP-specific T cells can induce dominant tolerance and reduce diabetes incidence

To measure the functional impact of antigen exposure on IGRP-specific cells, we studied diabetes incidence in NOD control and NOD-TII mice. There was no difference in diabetes between transgenic NOD mice that never expressed IGRP or expressed IGRP throughout life (Fig 6A, red vs white symbols) consistent with our previous studies indicating that deletion of IGRP-specific cells had no impact on diabetes (20, 25). Strikingly, however, induction of IGRP expression from week 10 onwards led to significant protection from diabetes (Fig 6A, red vs orange symbols). This observation suggested that exhausted IGRP-specific T cells can inhibit the function of T cells specific for other beta cell antigens. Indeed, it has been shown that exhausted T cells are not only hypofunctional but can also upregulate molecules that suppress local T cell immunity. Exhausted T cells upregulate CD39 and IL-10 which are both important molecules utilised by regulatory T cells (Tregs) to mediate suppressive functions(31, 32). Consistent with this idea, IGRP-specific cells in NOD-TII mice expressed high levels of IL-10 (Fig 6B) and CD39 (Fig 6C and D), while there was no difference in the frequency of FoxP3^+^ Tregs between NOD and NOD-TII mice (Fig 6E).

**Figure 6:**
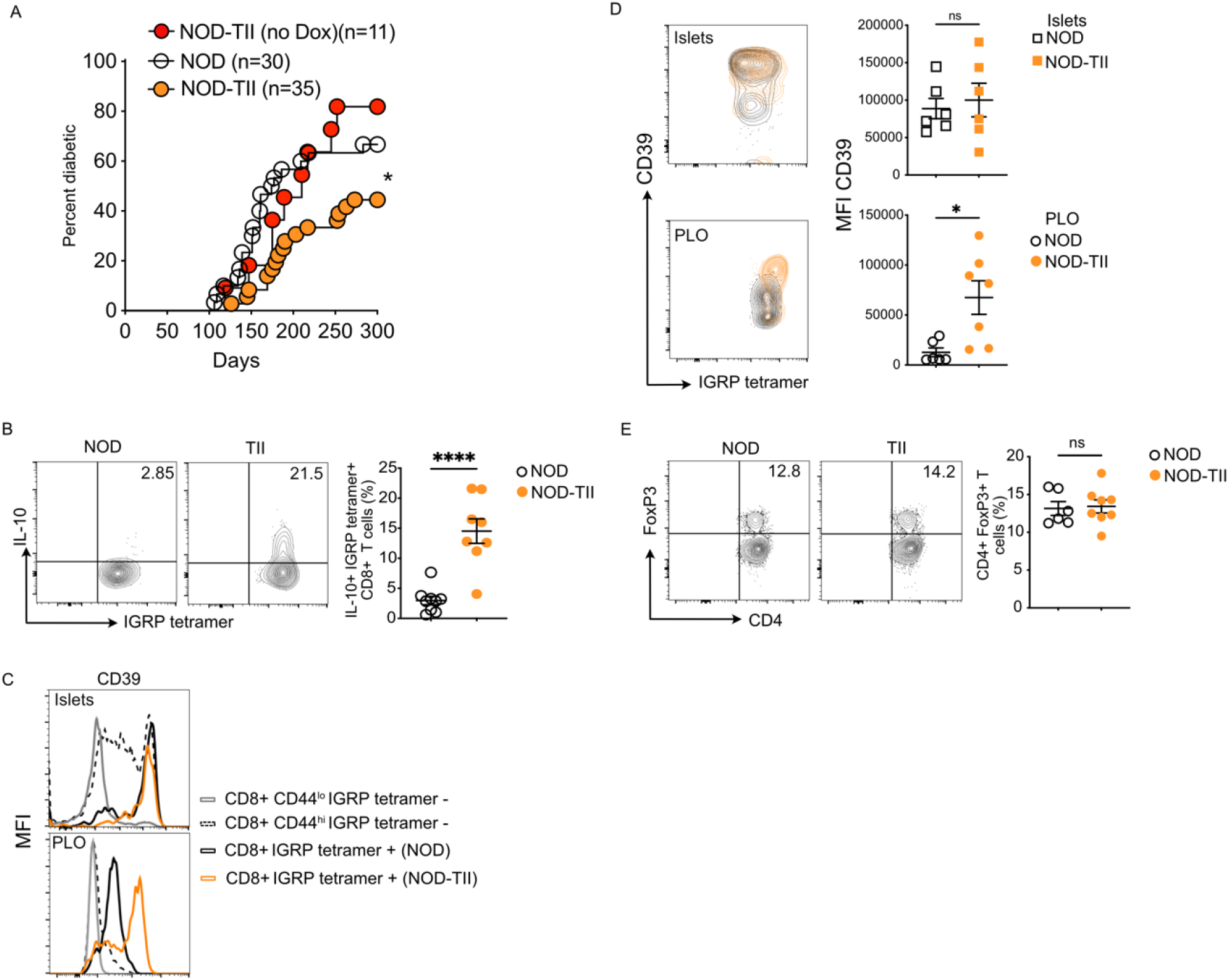
Exhausted IGRP-specific T cells induce dominant tolerance and reduce diabetes incidence. (A) Incidence of spontaneous diabetes until 300 days of age in cohorts of female NOD mice, NOD-TII mice expressing IGRP from birth and NOD-TII mice expressing IGRP from 10 weeks of age. Numbers in parentheses indicate the number mice analyzed. (B) Representative FACS plots and frequency of CD4^+^FoxP3^+^ Treg cells in PLO of 18–20-week-old NOD and NOD-TII mice. Numbers in FACS plots are frequency of Treg cells. (C) Representative FACS plots and frequency of IL-10 expressing IGRP-specific T cells enriched from PLO of 18–20-week-old NOD and NOD-TII mice and analyzed after re-stimulation with PMA and ionomycin for 4 hours. Numbers in FACS plots are frequency of IL-10 expressing cells. (D) Representative histograms showing expression of CD39 in the indicated CD8+ T cell subsets in islets (top panel) and PLO (bottom panel) of NOD and NOD-TII mice. Non-tetramer binding CD8+ T cell subsets shown are from NOD mice. (E) Mean fluorescence intensity quantification of CD39 in IGRP-specific T cells from islets (top) and PLO (bottom) of NOD and NOD-TII mice. Data pooled from 2-3 independent experiments. Statistical analysis performed using unpaired t-test (B,C,E). Survival curves in (A) compared using log-rank (Mantel-Cox) test. ns= not significant, * p<0.05, ****p<0.0001.

## Discussion

T cells express exhaustion-associated inhibitory receptors due to chronic antigen exposure in the islets. However, the islet microenvironment prevents differentiation to terminal exhaustion and the T cells are maintained in a less differentiated precursor of exhausted state with retained self-renewal features. These exhausted T cells undergo partial phenotypic and functional reversal upon antigen withdrawal when they leave the islets. The antigen experienced islet reactive T cells undergo differentiation to the terminally exhausted state when exposed to antigen in the extra islet microenvironment. Importantly, inducing differentiation to the terminally exhausted state in T cells specific for a single antigen led to protection from diabetes.

Our results are different to a recent study (23) showing that pancreas-infiltrating IGRP-specific T cells in NOD mice are short-lived effector T cells and do not display a transcriptional or phenotypic TOX driven exhaustion program. We used tetramer staining of T cells isolated from purified islets and found IGRP-specific T cells uniformly expressed high levels of TOX (Fig 1C), a critical transcription factor for the induction of CD8^+^ T cell exhaustion (33). The difference in the findings may in part reflect the use of different methods to isolate and analyze islet infiltrating T cells. We first isolate islets from the pancreas and then disperse them to single cells rather than the method used by Gearty to disperse the whole pancreas. This may result in non-islet and blood T cells being included in the analysis.

The predominance of T_PEX_ cells in the islets contrasts with what is seen in tumors and chronic viral infection (8, 34). In chronic viral infection, the T_PEX_ cells were mainly present in lymphoid tissues but were rare in the non-lymphoid tissues whereas the terminally exhausted T cells localized to both lymphoid and non-lymphoid tissues. T cells exposed to hypoxia due to high oxygen consumption by proliferating tumor cells may contribute to terminal exhaustion of T cells in the tumor microenvironment. Islet inflammation in NOD mice leads to development of tertiary lymphoid structures (35) and it is possible these structures support the development of T_PEX_ cells in NOD mice. Cancers have also been shown to develop intra tumoral tertiary lymphoid structures through chronic inflammatory signals. The presence of tertiary lymphoid structures in tumors has been linked to higher CD8^+^ T cell infiltration and improved responsiveness to immunotherapy, suggesting that tertiary lymphoid structures in the tumor support development of T_PEX_ cells(36). Interestingly, our scRNAseq data show high expression of *Ccr7* in T_PEX_ cells in the islets of NOD mice (Fig 3D). CCR7 is normally expressed in T cells in lymph nodes (e.g., central memory T cells) as compared to T cells in blood (e.g., effector memory T cells) or tissue (e.g., resident memory T cells).

The proportion of terminally exhausted IGRP-specific T cells does not change in NOD mice between 10 and 20 weeks of age in the islets but there are more in older NOD-TII mice than NOD mice (Fig 2). Islet specific T cells migrate between islet and peripheral lymphoid organs during the progression of diabetes meaning that the terminally exhausted, less functional IGRP-specific T cells that develop in the peripheral lymphoid organs continuously migrate into the islets of NOD-TII mice. This migration is important for progression of insulitis (35, 37). It is possible that these IL-10 secreting terminally exhausted IGRP-specific T cells protected NOD-TII mice from diabetes. Previous studies have shown that antigen delivery via nanoparticles coated with IGRP peptide bound to either MHC class 1 or class II molecules protected NOD mice from diabetes by inducing IL-10 secreting T cells (38, 39).

Our data indicate that NOD mice can be protected from diabetes at a time when an immune response is established against multiple antigens, by inducing exhaustion in T cells specific for a single antigen, IGRP. This suggests a dominant tolerance mechanism i.e. that the tolerant IGRP-specific T cells can inhibit T cells specific for other islet antigens. Terminally exhausted T cells are not only hypofunctional but also upregulate of molecules that suppress local T cell immunity such as CD39 and IL-10. Both are important molecules utilised by Tregs for suppressive functions. *CD39* (*Entpd1*) is expressed in terminally exhausted T cells (40) and acts via catabolising ATP and ADP to AMP and further degradation to adenosine via CD73. A recent study has shown that CD39 expressing terminally exhausted cells have suppressive capacity similar to conventional FoxP3 expressing CD4+ regulatory T cells(31). Several reports have identified that IL-10 secreting CD8^+^ T cells play a regulatory role in protecting against autoimmune disease (32, 41, 42). Conversely, blocking of IL-10 signaling improved the function of exhausted T cells in chronic viral infections (43) and simultaneous blockade of IL-10 and PD-1 pathways resulted in elimination of persistent viral infection (44).

We observed that IGRP-specific T cells partially lose their exhaustion-associated inhibitory markers as they traffic away from the site of antigen. This is similar to exhausted hepatitis C virus (HCV)-specific CD8^+^ T cells from infected patients recovering memory like features and some phenotypic reversal of exhaustion after elimination of virus with direct-acting antiviral therapy, and in chronic LCMV and CAR T cell models of exhaustion (45). Analysis of TCR diversity of HCV specific T cells following HCV cure suggests that transcriptional and epigenetic reprogramming may play a role in reversal of the exhaustion-associated phenotype (45). Future studies will need to determine whether the recovery of phenotypic exhaustion and memory like features that we observed following antigen removal are due to dedifferentiation of terminally exhausted T cells or exclusively due to outgrowth of new progeny from the T_PEX_ cell population.

The exhaustion program of T cells seems to be beneficial in restraining T cells in autoimmunity in some situations. Examples include patients with thyroid or islet antibodies who do not progress to clinical disease. These individuals rapidly develop autoimmune disease following checkpoint inhibitor treatment (17, 46). Maintenance of the ability to destroy target tissue despite chronic antigen exposure occurs in autoimmunity and by definition T cell exhaustion as a tolerance mechanism is ineffective in people with autoimmune diseases. In tumors, there is a progressive increase in exposure to antigen as the tumor grows, and the T cells progressively become more exhausted with attrition of T_PEX_ cells (47). Studies in the LCMV model of T cell exhaustion showed that an increased amount of antigen can drive more severe T cell exhaustion. In the immune response against the beta cells, the amount of target tissue is fixed or declining (48), unlike in disease such as cancer and viral infections in which the antigens have the capacity to expand. The T cells are exposed to a fixed and limited amount of transgenic antigen in our model. Modulating the amount of antigen in the peripheral lymphoid organs could have an impact on the proportion of precursor versus terminally exhausted T cells.

In summary, we found that T cells undergo an exhaustion program in NOD mice. However, this process is not complete as these cells persist mainly as precursors of exhausted T cells. Antigen exposure in the extra-islet environment induced terminal differentiation of T cells. Terminally exhausted T cells specific for a single antigen led to protection from diabetes. The predominance of T_PEX_ cells in the islets implies that factors within the islet support them and their elimination from that site is the highest priority.

## Methods

### Mice

NOD/Lt mice were bred and housed at the Bioresources Centre, St. Vincent’s Hospital, Fitzroy. NOD-IEα-tTA mice that drive the expression of tetracycline transactivator (tTA) under control of the MHC class II IEα promoter have been previously described(49) and were obtained from Prof. C. Benoist and Prof. D. Mathis (Dept of Pathology, Harvard Medical School, Boston, Massachusetts, USA). TetO-IGRP transgenic mice were crossed with NOD-IEα-tTA mice to generate dual transgenic NOD-TII mice as previously described(25). NOD8.3 mice, expressing the TCRαβ rearrangements of the H-2Kd-restricted, β cell-reactive, CD8+ T cell clone NY8.3, have been previously described(50). All mice were bred, maintained and used under specific pathogen free conditions at St Vincent’s Institute (Melbourne, Australia). All experimental procedures followed the guidelines approved by the institutional animal ethics committee.

### Doxycycline treatment

Untreated NOD-TII mice constitutively express IGRP in antigen presenting cells (APCs). To turn-off IGRP expression, doxycycline hyclate (Dox) (Sigma-Aldrich) was administered via drinking water at a concentration of 10 mg/L. Water bottles were changed thrice weekly. NOD-TII mice receiving water with Dox were then given unmedicated water to re-express IGRP.

### Real-time polymerase chain reaction (qPCR)

For total RNA extraction, spleens were harvested in cold Phosphate Buffered Saline (PBS). Tissue homogenates were prepared in RNA lysis buffer RLY1 (Bioline) from a 15 mg slice of tissue using a tissue homogenizer. RNA was isolated using ISOLATE II RNA Mini Kit (Bioline) according to the manufacturer’s instructions. Eluted RNA was further purified with TURBO DNA-free™ Kit (Invitrogen). A NanoDrop 2000 UV-Vis Spectrophotometer was used to verify RNA content. Complementary DNA synthesis was performed using High Capacity cDNA Reverse Transcription Kit with RNase Inhibitor (Applied Biosystems). AmpliTaq Gold Taqman Assay (Applied Biosystems) was performed in a Roche LightCycler 480 Instrument II. Taqman gene expression probes used in the study are listed in supplementary table S2. Relative expression of genes of interest was determined by normalization to indicated reference genes using differential threshold cycle (ΔCT, 2^−ΔΔC_*T*_^) analysis.

### CFSE labelling and adoptive transfer

Single cell suspensions of NOD 8.3 splenocytes were prepared and labelled with carboxyfluorescein succinimidyl ester (CFSE, Thermo Fisher) as previously described(20). 5 x 10^6^ CFSE labelled cells were injected intravenously into tail veins of NOD-TII mice with or without induced IGRP expression. Hosts were sacrificed after 5 days and their inguinal lymph nodes (ILN), pancreatic lymph nodes (PLN) and pancreatic islets were examined for CFSE+ cells by flow cytometry.

### Islet isolation

Islets were isolated using collagenase P (Roche, Basel, Switzerland) and Histopaque-1077 density gradients (Sigma-Aldrich) as previously described(37). Islets were dissociated via bovine trypsin (Calbiochem) hydrolysis and single cell suspensions were processed for flow cytometry.

### Flow Cytometry

Single cell suspensions were prepared from pooled peripheral lymphoid organs (spleen and non-draining lymph nodes, PLO) and islets. Cell suspensions from PLO were treated with ammonium chloride buffer to lyse red blood cells. For surface staining, cells were stained with fluorescently labelled antibodies on ice for 30 minutes. All antibodies used are listed in Supplementary table S2. Propidium Iodide (PI, Calbiochem) was added prior to flow-cytometry analysis to exclude dead cells. IFNγ and IL-10 were detected intracellularly using the Cytofix/Cytoperm Kit (BD Biosciences) following incubation with PMA and Ionomycin for 4 hours at 37°C. The list of antibodies used for flow cytometry is provided in supplementary table S1. Intracellular staining for TOX, TCF-1 and FoxP3 was performed using FoxP3/Transcription Fixation/Permeabilization kit (eBiosciences). Data were collected with a LSR Fortessa (Becton Dickinson) or Cytek Aurora (Millenium Biosciences) flow cytometer and analyzed with FlowJo (v10.8.1) (Tree Star, Ashland, USA) software.

### Tetramer staining and magnetic bead-based enrichment

The tetramer and magnetic bead-based enrichment assay has been previously described(26). Briefly, single cell suspensions from PLO (pooled spleen and non-draining lymph nodes comprising of 2 inguinal and 4 mesenteric lymph nodes per sample) were stained with phycoerythrin (PE)-conjugated IGRP_204-216_ (VYLKTNVFL) H2-Kd tetramer (ImmunoID, Parkville, Victoria, Australia), for 1 hour on ice, washed and incubated with anti-PE magnetic beads (Miltenyi Biotec, Cologne, Germany) followed by magnetic separation using an AutoMACSpro (Miltenyi Biotec) according to manufacturer’s instructions. The tetramer enriched fractions were stained with cell-surface markers and analyzed by flow cytometry. Gating strategy for tetramer enrichment was as follows: single cells were gated on forward and side scatter, and dead cells excluded using propidium iodide. From the live cell population, CD3^+^ dump^-^ (dump= CD11c, CD11b, B220 and F4/80) cells were gated as the T cell population for analysis of tetramer binding CD4^-^ CD8^+^ T cells (Supplementary Fig S2).

### Cell sorting, library preparation and sequencing for sc-RNAseq

For sc-RNAseq analysis, live CD45^+^ cells were FACS sorted from dispersed islets pooled from 2-3 NOD mice (15-16 weeks of age) per sample. Sorted CD45^+^ cells were washed and resuspended in RPMI cell culture medium (Gibco) containing 10% FCS at a density of 1200 cells/μl and loaded on to Chromium Controller (10X Genomics). 3 samples from 2 independent experiments were processed further using Chromium Single cell 3’ Gel bead kit (v 3.1) and library construction kit as per manufacturer’s instructions. The libraries were quantified using the Agilent Bioanalyzer High sensitivity Chip and sequenced on the Illumina NovaSeq PE150 platform (Novogene AIT Genomics, Singapore).

### Single-cell RNAseq dataset alignment and clustering

Sequencing files were demultiplexed and aligned to *Mus musculus* transcriptome reference (mm10), and count matrices were extracted using Cell Ranger software v6.0.1 (10X Genomics). The expression matrices were then imported into R (v4.0.4) and processed with Seurat (v4.0.0). Following cell-cycle regression, doublet removal and filtering of cells based on counts, genes and percentage of mitochondrial genes, the remaining cells in each sample were normalized and integrated by SCTransform to correct for batch effects. Dimensionality reduction was performed by unsupervised principal component analysis (PCA) and uniform manifold approximation and projection (UMAP) embedding for each sample. 22,616 CD45^+^ cells from 3 samples were re-clustered using a K-nearest neighbor (KNN) graph followed by Louvain clustering using 12 PCs and a resolution of 1.2. The resulting cell clusters were visualized using t-distributed stochastic neighbor embedding (t-SNE) and UMAP. Cell-type meta-clusters were annotated using a list of pre-defined marker genes and the FindMarkers function to identify cell-type specific gene signatures within each cluster (Supplementary figure X). A subset of 16,585 cells identified as T-cells were re-clustered using 14 PCs and a resolution of 1.5. Cd8a and Cd4 expression was used to identify islet CD8^+^ and CD4^+^ T-cells. Finally, 4,520 CD8^+^ T-cells were re-clustered using 14 PCs and a resolution of 1.2 and a pre-determined set of genes was used to annotate CD8^+^ T-cell subsets. PCA cut-offs were established by determining the number of PCs accounting for 90% of the variance.

### Diabetes incidence

Female mice were monitored for spontaneous diabetes over a 300-day time-course. Diabetes onset was monitored by weekly measurement of urine glucose levels using Diastix (Bayer Diagnostics). Blood glucose levels were measured in mice with glycosuria (>110 mmol/L) using Advantage II Glucose strips (Roche). Animals displaying two consecutive blood glucose measurements of ≥ 15mmol/L were considered diabetic.

### Quantification and Statistical analysis

All statistical analyses were performed using GraphPad Prism 9 software (GraphPad, San Diego, USA). A two-tailed unpaired Student’s t-test was used for comparisons between two groups. Multiple comparisons were performed using one-way ANOVA with Tukeys post-hoc test. Diabetes incidence curves were compared using Log-rank (Mantel-Cox) test. In all graphs, each symbol represents an individual sample, and the error bars represent the mean ± standard error of mean. *p < 0.05, **p < 0.01, ***p < 0.001.

## Supporting information

Supplementary Table S1-antibodies

Supplementary Table S2-qPCR probes

## Author contributions

CS, GJ, CTJK, MKC, DdG, TG, PT and EP performed experiments, XL analyzed the data, CS, GJ, DdG, CTJK and AK analyzed the data and wrote the manuscript, HET, BK and TWHK designed the study, analyzed the data and wrote the manuscript. BK and TWHK supervised the study.

## Acknowledgments

The authors thank T. Catterall, S. Fynch, E. Batleska, (St. Vincent’s Institute) V. Moshovakis, R. Greaves and E. Gumbrell (St. Vincent’s Hospital) for excellent technical assistance and animal husbandry. This work was funded by a National Health and Medical Research Council of Australia (NHMRC) Program grant (GNT1150425). St Vincent’s Institute receives support from the Operational Infrastructure Support Scheme of the Government of Victoria.

