## Supplementary Figures and legend for "Extra-islet expression of islet antigen boosts T-cell exhaustion to prevent autoimmune diabetes"

**Supplementary Figure legend**

**Supplementary Figure 1: sc-RNAseq analysis and clustering of immune cells in the islets of NOD mice**

(A) Uniform Manifold Approximation and Projection (UMAP) plot of islet infiltrating CD45+ cells in 15-16 weeks old NOD mice (B) Expression of canonical immune cell markers in clusters of islet infiltrating CD45+ cells in A. (C) UMAP plot of re-clustered T cells from panel A. (D) Expression of canonical T cell markers in re-clustered T cells. T cells expressing CD4 were excluded for further analysis.

**Supplementary Figure 2: Analysis of immune cell population in islets and peripheral lymphoid organs of NOD and NOD-TII mice.**

(A)Representative plots showing gating strategy for analysis of IGRP -specific T cells from peripheral lymphoid organs and (B) islets of NOD and TII mice. Cells were gated on forward scatter (FSC) and side scatter (SSC), propidium iodide– (PI) live cells, CD3+, CD4- CD8+ and tetramer. Representative staining for indicated markers is shown. (C) Representative plots showing gating strategy for analysis of co-stimulatory markers on antigen presenting cells (APCs) from spleen in NOD and TII mice. Cells were gated on forward scatter (FSC) and side scatter (SSC), propidium iodide– (PI) live cells, CD19+ B cells and CD3+ T cells were excluded and CD19- CD3- cells were further classified into MHC II + CD11c+ dendritic cells. Representative plot showing MFI of indicated marker is shown. (D) Representative flow cytometry plots showing gating strategy to sort islet infiltrating CD45+ immune cells for sc-RNAseq analysis. Cells were gated on forward scatter (FSC) and side scatter (SSC), propidium iodide– (PI) live cells.CD45 + cells and processed for sc-RNAseq analysis.

**Supplementary Figure 3: Analysis of exhaustion related markers and co-stimulatory receptors on antigen-presenting cells in NOD and NOD-TII mice.**

(A) Representative FACS plots showing co-expression of TCF1 and Slamf6 on IGRP- specific T cells in islets and PLO of NOD mice. Numbers are the frequency of cells in each quadrant. (B) Proportion of Slamf6+ and TIM3+ IGRP specific CD8+ T cells in PLN of 16-18-week-old NOD mice. (C) Representative histogram showing expression of PD-1 on $CD8^+$ T cell subsets. (D) Quantitative RT-PCR for G6pc2 in splenic lysates of wild-type NOD and NOD-TII mice with no induction of IGRP expression. Data show dCT values (mean±SEM) of individual mice from 2 independent experiments. (E) Expression of MHC II, CD80, CD86 and CD40 was examined on $CD3^-$ $CD19^-$ MHC class II $^+$ $CD11c^+$ antigen presenting cells (APCs) in dispersed islets and pooled peripheral lymphoid organs (PLO). Mean fluorescence intensity (MFI) quantification shown in NOD (top panel) and TII mice (bottom panel). NOD.$Rag1^{-/-}$ islets were transplanted under the kidney capsule of NOD and NOD-TII mice. (F) Enumeration of IGRP-specific T cells and (G) percentage of $IFN\gamma^+$ IGRP- specific T cells after restimulation with PMA and ionomycin, enriched from PLO of NOD, NOD-TII and NOD-TII mice with transient IGRP expression. Each symbol in the scatter plots (mean±SEM) represents data from an individual mouse. Data representative of 2 independent experiments. Statistical analyses performed using unpaired t-test (D, E) and One-way ANOVA with Sidaks post-test (G,H) ns= Not significant, * $p<0.05$, ** $p<0.01$, **** $p<0.0001$.

**Supplementary Figure S1**

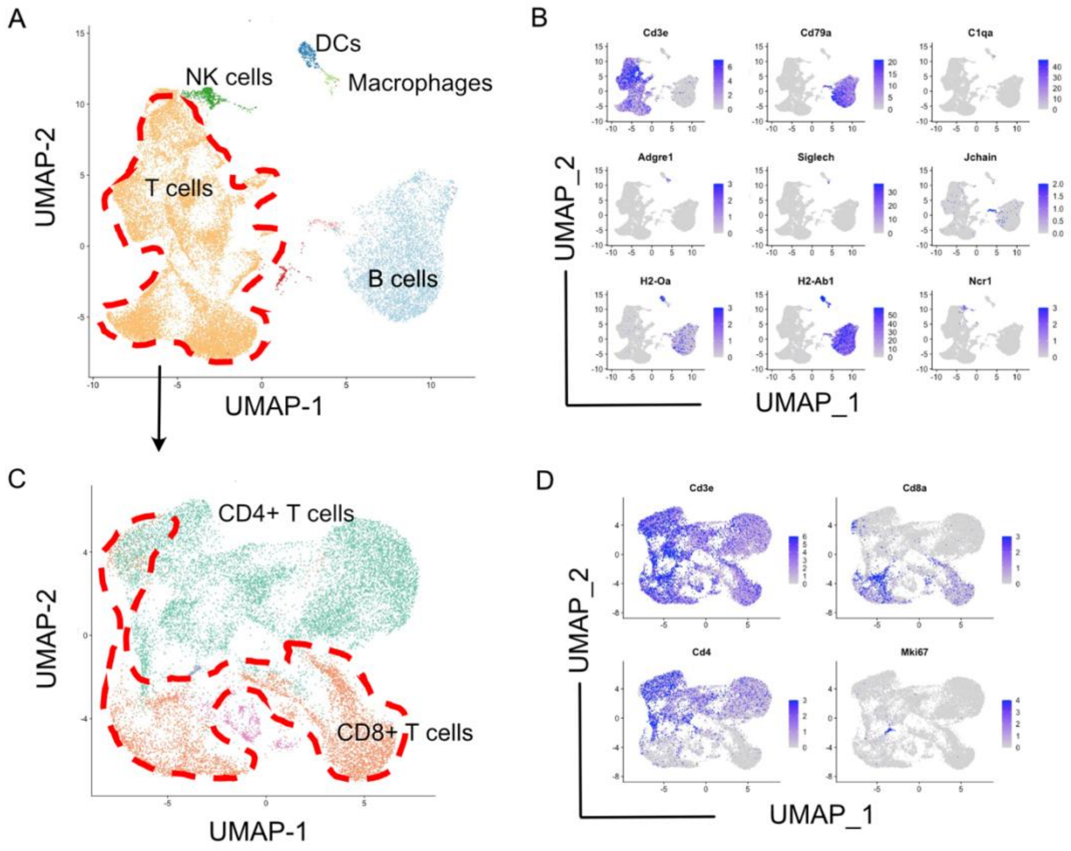

**Supplementary Figure S2**

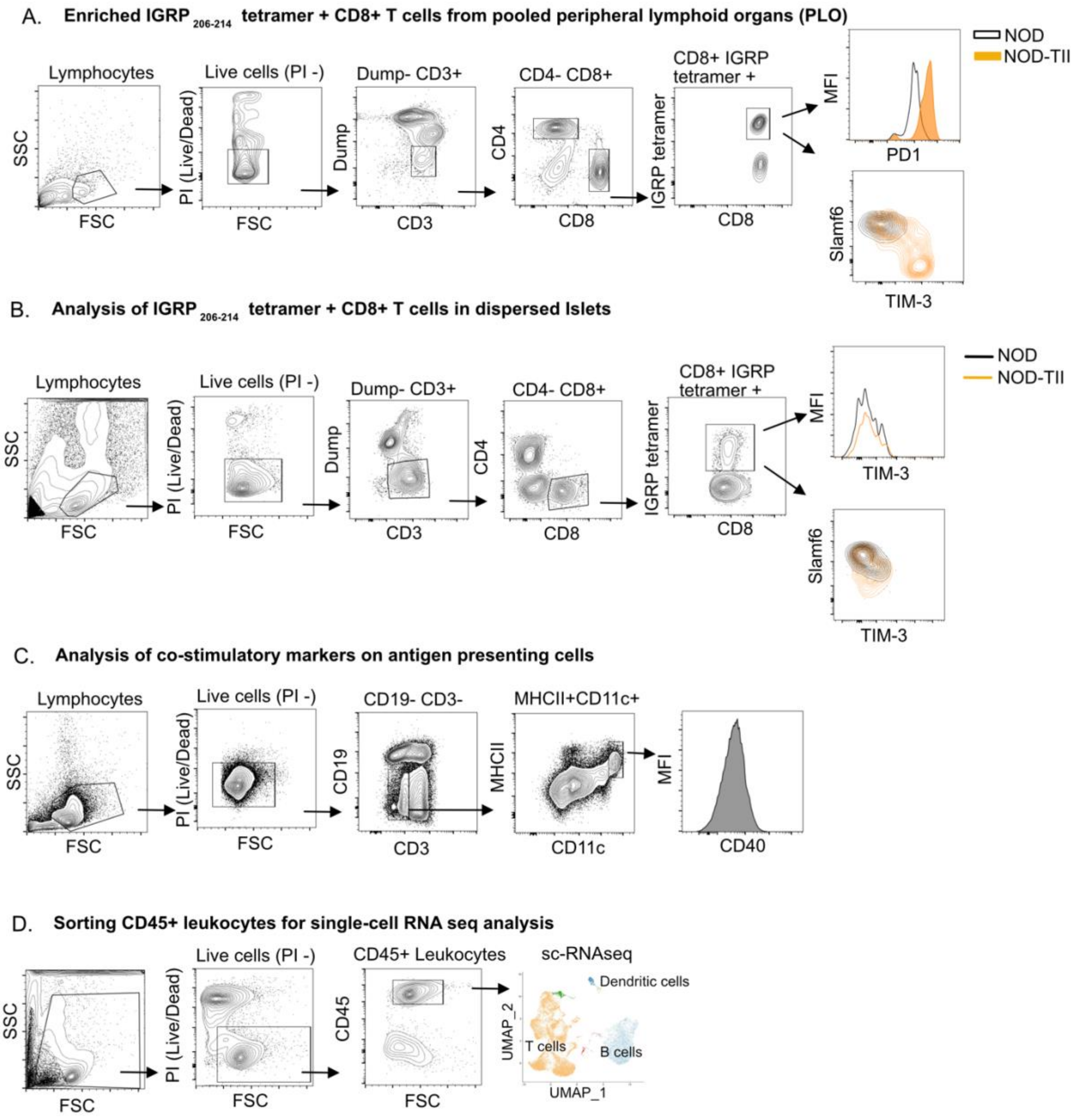

**Supplementary Figure S3**

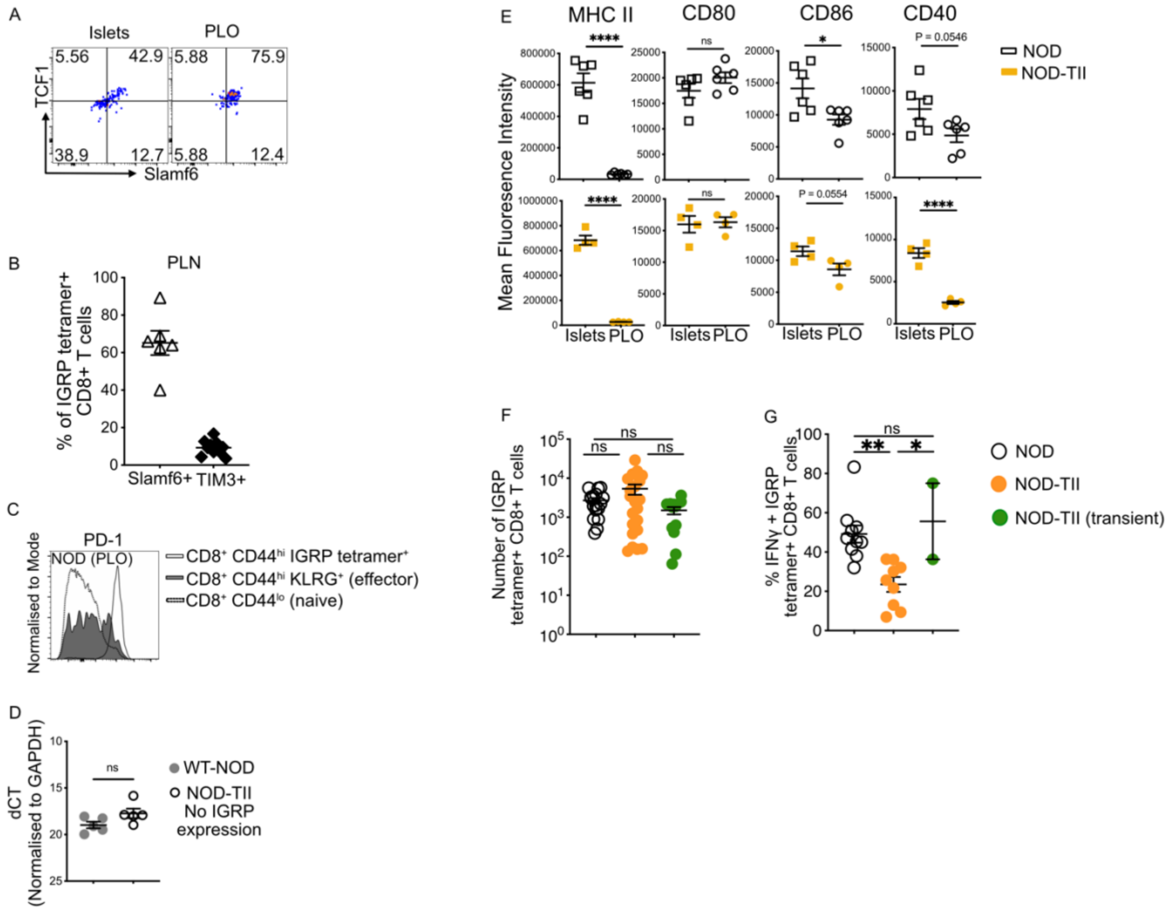
