## Supplementary Table S1-antibodies for "Extra-islet expression of islet antigen boosts T-cell exhaustion to prevent autoimmune diabetes"

**Supplementary Table S1: List of Antibodies**

| Target. | Supplier. | Catalog ID. |
|---|---|---|
| CD3 (500A2) V500 | BD Biosciences | 560771(RRID:AB_1937314) |
| CD8 (53-6.7) APC | BioLegend | 100712 (RRID:AB_312751) |
| CD8 (53-6.7) PercpCy5.5 | BD Biosciences | 551162 (RRID:AB_394081) |
| CD8 (53-6.7) BV711 | Biolegend | 100759(RRID:AB_2563510) |
| CD4 (GK1.5) APC-cy7 | BioLegend | 100414 (RRID:AB_312699) |
| CD4 (GK1.5) FITC | BDBiosciences | 553729 (RRID:AB_395013) |
| CD4 (GK1.5) BV650 | BDBiosciences | 563232 (RRID:AB_2738083) |
| CD45 (30-F11) PerCP-Cy5.5 | BioLegend | 103132(RRID:AB_893340) |
| CD45 (30-F11) PE-cy7 | BDBiosciences | 552848(RRID:AB_394489) |
| CD44 (1M7) PE-Cy7 | BioLegend | 103030 (RRID:AB_830787) |
| PD-1 (29f.1A12) BV605 | BioLegend | 135220 (RRID:AB_2562616) |
| TIM-3 (215008) AF488 | RD Systems | FAB1529G (RRID:NA) |
| TIM-3 (215008) AF700 | RD Systems | FAB1529N (RRID:NA) |
| TIGIT (VSIG9) PE/Dazzle | BioLegend | 142110 (RRID:AB_2566573) |
| Tox (TXRX10)-eFlour660 | eBioscience | 50-650282 (RRID:AB_2574265) |
| Slamf6(REA109)-FITC | Miltenyi | 130-118-597 (RRID:AB_2733606) |
| CD11b (M1/70) eFluor450 | eBioscience | 48011282(RRID:AB_1582236) |
| CD80 (16-10A1)-PE-cy7 | Biolegend | 104733(RRID:AB_2563112) |
| CD86 (GL-1)-BV605 | BDBiosciences | 563055(RRID:AB_2737977) |
| CD40 (1C10)-PE | Biolegend | 102805(RRID:NA) |
| CD19(6D5)-APC-cy7 | Biolegend | 115529(RRID:AB_830706) |
| MHC class II (OX-6)-FITC | BDBiosciences | 554928(RRID:AB_395605) |
| KLRG1 (2F1)-APC | Biolegend | 138412(RRID:AB_10641560) |
| CD11c (N418) eFluor450 | eBioscience | 48011482 (RRID:AB_1548654) |
| B220 (RA3-6B2) eFluor450 | eBioscience | 48045282 (RRID:AB_1548761) |
| B220 (RA3-6B2) PE | BioLegend | 103208 (RRID:AB_312993) |
| Ly6G (RB6-8C5) eFluor450 | eBioscience | 48-5931-82 (RRID:AB_1548788) |
| F4/80 (BM8) eFlour450 | eBioscience | 48-4801-82 (RRID:AB_1548747) |
| CD16/CD32 (2.4G2) | BDBiosciences | 553142(RRID:AB_394656) |
| IFNG (XMG1.2)-FITC | eBioscience | 17-7311-82 (RRID:AB_469504) |
| IL-10 (JES5-16E3)-APC | eBioscience | 17-7101-81 (RRID:AB_ 469501) |
| FoxP3 (FJK-16s) APC | eBioscience | 17-5773-82 (RRID:AB_469457) |
| FoxP3 (FJK-16s) PE | eBioscience | 12-5773-82 (RRID:AB_465936) |
| MHC-1 (SF1-1.1) Biotin | BD Biosciences | 553564 (RRID:AB_394922) |
| Streptavidin APC | BioLegend | 405207 (RRID:NA) |
