## Supplementary Table S2-qPCR probes for "Extra-islet expression of islet antigen boosts T-cell exhaustion to prevent autoimmune diabetes"

**Supplementary Table S2: List of qRT-PCR probes**

| Protein name | Mouse gene name | Taqman assay |
|---|---|---|
| PD-1 | PDCD1 | Mm01285676_m1 |
| TIM3 | HAVCR2 | Mm00454540_m1 |
| TIGIT | TIGIT | Mm03807522_m1 |
| CD38 | CD38 | Mm00483143_m1 |
| EOMES | EOMES | Mm01351984_m1 |
| T-Bet | TBX21 | Mm00450960_m1 |
| PR Domain 1 | PRDM1 | Mm00476128_m1 |
| C-MAF | MAF | Mm02581355_s1 |
| Lag3 | Lag3 | Mm00493071_m1 |
| Tox | TOX | Mm00455231_m1 |
| IGRP | G6PC2 | Mm00491176_m1 |
| Beta actin | ACTB | Mm00607939_s1 |
| 18s RNA | Rn18s | Mm03928990_g1 |
| GAPDH | Gapdh | Mm99999915_g1 |
